# Improving the efficiency of de Bruijn graph construction using compact universal hitting sets

**DOI:** 10.1101/2020.11.08.373050

**Authors:** Yael Ben-Ari, Dan Flomin, Lianrong Pu, Yaron Orenstein, Ron Shamir

**Affiliations:** Blavatnik School of Computer Science, Tel-Aviv University, Tel-Aviv, Israel; School of Electrical and Computer Engineering, Ben-Gurion University of the Negev, Beer-Sheva, Israel

## Abstract

High-throughput sequencing techniques generate large volumes of DNA sequencing data at ultra-fast speed and extremely low cost. As a consequence, sequencing techniques have become ubiquitous in biomedical research and are used in hundreds of genomic applications. Efficient data structures and algorithms have been developed to handle the large datasets produced by these techniques. The prevailing method to index DNA sequences in those data structures and algorithms is by using *k*-mers (*k*-long substrings) known as minimizers. Minimizers are the smallest *k*-mers selected in every consecutive window of a fixed length in a sequence, where the smallest is determined according to a predefined order, e.g., lexicographic. Recently, a new *k*-mer order based on a universal hitting set (UHS) was suggested. While several studies have shown that orders based on a small UHS have improved properties, the utility of using them in high-throughput sequencing analysis tasks has not been demonstrated to date.

Here, we demonstrate the practical benefit of UHSs for the first time, in the genome assembly task. Reconstructing a genome from billions of short reads is a fundamental task in high-throughput sequencing analyses. De Bruijn graph construction is a key step in genome assembly, which often requires very large amounts of memory and long computation time. A critical bottleneck lies in the partitioning of DNA sequences into bins. The sequences in each bin are assembled separately, and the final de Bruijn graph is constructed by merging the bin-specific subgraphs. We incorporated a UHS-based order in the bin partition step of the Minimum Substring Partitioning algorithm of Li *et al*. (2013). Using a UHS-based order instead of lexicographic- or random-ordered minimizers produced lower density minimizers with more balanced bin partitioning, which led to a reduction in both runtime and memory usage.

## 1 Introduction

Large amounts of DNA sequencing data are generated today in almost any biological or clinical study. Due to the low cost of sequencing, it has become standard to probe and measure molecular interactions and biomarkers using DNA read quantities [16]. Technologies based on high-throughput sequencing (HTS) have been developed for the major genomics tasks: genetic and structural variation detection, gene expression quantification, epigenomic signal quantification, protein binding measurements, and many more [6]. A first step in utilizing all these data types is the computational analysis of HTS data. Key challenges include read mapping to a reference genome, read compression, storing reads in a data structure for fast querying, and finding read overlaps. As a result, many computational methods were developed to analyze HTS data, and the development of new methods is ongoing [1].

Many methods for analyzing HTS data use minimizers to obtain speed-up and reduce memory usage [17, 18, 8]. Given integers *w* and *k*, the *minimizer* of an *L* = *w* + *k* − 1-long sequence is the smallest *k*-mer among the *w* contiguous *k*-mers in it, where the smallest is determined based on a predefined order, e.g., lexicographic [19]. For a longer sequence, all *L*-long windows are scanned and the minimizer is selected in each one (Figure 1a). Using the minimizers to represent the *L*-long windows has three key advantages: (i) the sampling interval is small; (ii) the same *k*-mers are often selected from overlapping windows; and (iii) identical windows have the same minimizer. Minimizers help design algorithms that are more efficient in both runtime and memory usage by reducing the amount of information that is processed while losing little information. Minimizers were shown to be helpful and are used in many different settings, such as partitioning input sequences [4, 17, 18], generating sparse data structures [5, 21], and sequence classification [20].

**Figure 1:**
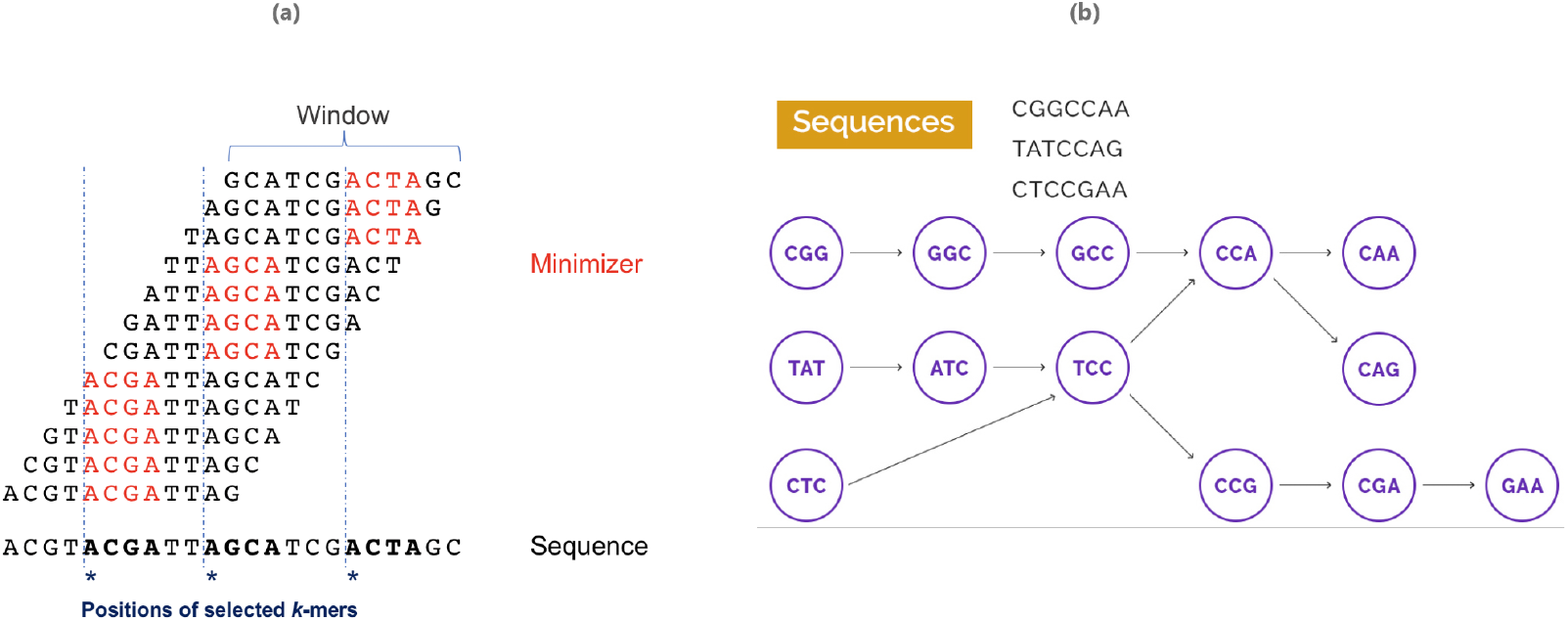
Illustrations of preliminary definitions. (a) A minimizers scheme (*k* = 4, *w* = 9). The input sequence is broken into windows of length *L* = *w* + *k* − 1 = 12, and the minimizer in each window is selected. Consecutive windows tend to select the same minimizer. The positions of the selected *k*-mers constitute a sampling of the original sequence. (b) de Bruijn graph of order 3 for three DNA sequences. The vertices are the 3-mers contained in the set of sequences. Edges connect two vertices if the 4-mer they represent is contained in a sequence in the set.

Recently, the concept of a universal hitting set (UHS) was introduced as a way to improve minimizers [15]. For integers *k* and *L*, a set of *k*-mers *U_kL_* is called a UHS if every possible sequence of length *L* contains at least one *k*-mer from *U_kL_* as a contiguous substring. It was shown that by using a UHS of small size, one can design an order for a minimizer scheme that results in fewer selected *k*-mers compared to the orders commonly used in current applications (i.e., lexicographic or random orders) [12]. Therefore, using UHSs has the potential to provide smaller signatures than currently used orders, and as a result reduce runtime and memory usage of sequencing applications. We and others recently developed algorithms to generate small UHSs [15, 12], but so far the prevailing methods in HTS analysis employ a lexicographic or random order. To date, no method has been developed to take advantage of the improved properties of UHSs.

In this study we demonstrate, for the first time, the practical benefit of UHSs on a HTS analysis task: de Bruijn graph construction for genome assembly by a disk-based partition method. We introduce a UHS into the graph construction step of the Minimum Substring Partition assembly algorithm [10]. The introduction of the UHS into the algorithm defines a new minimizers ordering, substantially changing the execution of all the steps of the algorithm, but producing exactly the same final output. In tests on several genomic datasets, the new method had lower memory usage, shorter runtime and more balanced disk partitions. The code of our method is publicly available at github.com/Shamir-Lab/MSP_UHS.

## 2 Background and Preliminaries

### 2.1 Definitions

#### 2.1.1 Basic definitions

A *read* is a string over the DNA alphabet Σ = {*A, C, G, T*}. A *k*-mer is a *k*-long string over Σ. Given a read *s*, |*s*| = *n*, *s*[*i, j*] denotes the substring of *s* from the *i*-th character to the *j*-th character, both inclusive. (Here and throughout, substrings are assumed to be contiguous.) *s* contains *n* − *k* + 1 *k*-mers: *s*[0, *k* − 1], *s*[1, *k*],…, *s*[*n* − *k*, *n* − 1]. Two *k*-mers in *s* that overlap in *k* − 1 letters, i.e., *s*[*i*, *k* + *i* − 1] and *s*[*i* + 1, *k* + *i*], are called *adjacent* in *s*.

#### 2.1.2 De Bruijn graphs

Given a set of strings *S* = {*S*_0_, *S*_1_, *S*_2_,…, *S*_*m−1*_} over Σ and an integer *k* ≥ 2, the *de Bruijn graph* of *S* of order *k* (Figure 1b) is a directed graph *dBG_k_*(*S*) = (*V, E*) where:

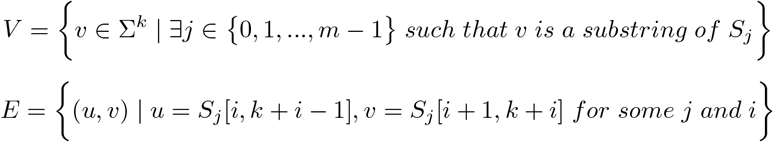

Modern genome assembly algorithms are based on de Bruijn graph construction. This process breaks each input read into *k*-mers (vertices in the graph) and then connects adjacent *k*-mers according to their overlap relations in the reads (edges). The graph represents the reconstructed genome. This process can assemble very large quantities (even billions) of reads. The most memory consuming and time-intensive part in assembly algorithms is the de Bruijn graph construction step [10].

#### 2.1.3 Minimizers and orders

An order *o* on Σ^*k*^ is a one-to-one function *o* : Σ^*k*^ → {1, 2,…, |Σ|^*k*^}. *k*-mer *m*_1_ is smaller than *k*-mer *m*_2_ according to order *o* if: *o*(*m*_1_) < *o*(*m*_2_). In other words, an order is a permutation on the set of all *k*-mers. A *minimizer* for a triplet (*s, o, k*) is the smallest *k*-long substring *m* in sequence *s* according to order *o*. We also call *m* the *o*-minimizer *k*-mer in *s*. A *minimizers scheme* is a function *f_k,w_* that selects the start position of a minimizer *k*-mer in every sequence of length *L* = *w* + *k* − 1, i.e., *f* : Σ^*w*+*k*−1^ → [0 : *w* − 1] (Figure 1a).

#### 2.1.4 Particular density

The set of selected positions of a scheme *f_k,w_* on a string *s* is *M_f__k,w_* (*s*) = {*i* + *f_k,w_*(*s*[*i*, *i* + *k* + *w* − 2]) where 0 ≤ *i* ≤ |*s*| − *w* − *k* + 1} (asterisks in Figure 1a). The *particular density* of a scheme *f_k,w_* on a string *s* is the proportion of *k*-mers selected:

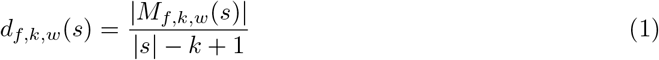

Particular density was used in previous works (e.g., [22]) as a measure of efficiency of the scheme on a particular sequence. The trivial upper and lower bounds for the density are 1/*w* ≤ *d_f,k,w_* ≤ 1, where 1/*w* corresponds to scanning the sequence from left to right and selecting exactly one position in every new non-overlapping window, and 1 corresponds to selecting every position [12]. In general, lower density can lead to greater computational efficiency and is therefore desirable.

#### 2.1.5 Universal hitting sets

A set of *k*-mers *M hits* sequence *s* if there exists a *k*-mer in *M* that is a substring in *s*. A *universal hitting set* (UHS) *U_kL_* is a set of *k*-mers that hits every *L*-long string over Σ. A trivial UHS always exists by taking all (|Σ|^*k*^) *k*-mers. A UHS *M* can be used in a minimizers scheme as follows: Define an order on *M*’s *k*-mers, and for any *L*-long window select the minimum *k*-mer from *M* in the window according to the defined order. The universality of *M* guarantees that there will always be at least one *k*-mer from *M* in any *L*-long window.

### 2.2 Minimum Substring Partitioning

The Minimum Substring Partitioning (MSP) method is a memory-efficient and fast algorithm for de Bruijn graph construction [10]. MSP breaks reads into multiple bins so that the *k*-mers in each bin can be loaded into memory, processed individually, and later merged with other bins to form the de Bruijn graph. The lexicographically smallest *k*-mer in each sequence window (i.e., the minimizer) is used as key for that window.

MSP partitions *L*-long windows into multiple disjoint bins, in a way that tends to retain adjacent *L*-mers in the same bin. This has two advantages: (i) consecutive *L*-mers are combined into *super L-mers* (substrings of length ≥ *L*), which reduces the space requirements; (ii) local assembly can be performed on the bins in parallel, and later all assemblies are merged to generate a global assembly. MSP is motivated by the fact that adjacent *L*-mers tend to share the same minimizer *k*-mer, since there is an overlap of length *L* − 1 between them. Figure 2 shows an example of the partitioning step of MSP with *L* = 10 and *k* = 3. In this example, the first four *L*-mers share the minimizer *AAC*; and the last four *L*-mers share the minimizer *AAA*. In this case, instead of generating all seven *L*-mers separately, MSP generates only two super *L*-mers. The first four *L*-mers are combined into *TGGCGAACGTAA*, and this super *L*-mer is assigned to the bin labeled *AAC*. Similarly, the last four *L*-mers are combined into a super *L*-mer *GAACCGTAAAGT*, and this super *L*-mer is assigned to the bin labeled *AAA*. In general, given a read *r* = *r*_0_*r*_1_… *r*_*n*−1_, if the *j* adjacent *L*-mers from *r*[*i, i + L* − 1] to *r*[*i* + *j* − 1, *i* + *j* + *L* − 2] share the same minimizer *m* (and *j* is maximal with regard to that property), then the super *L*-mer *r*_*i*_*r*_*i*+1_… *r*_*i+j+L−2*_ is assigned to the bin labeled *m* without breaking it into *j* individual *L*-mers. This procedure reduces memory usage as instead of keeping *j* · *L* characters in memory, only *j* + *L* − 1 characters are kept. If *j* tends to be large, this strategy dramatically reduces memory usage. To reduce the number of bins, MSP warps the bins using a hash function into a user-defined number of bins *b*.

**Figure 2:**
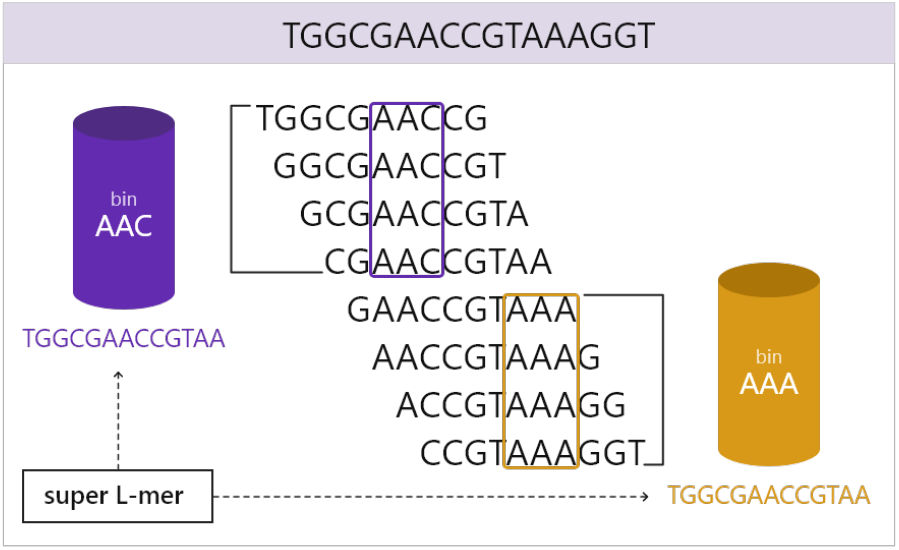
The partitioning step of the MSP method. A read is scanned in windows of length 10. The 3-mer minimizer in each window is marked with the rectangles.

Li *et al.* argued that the maximum number of distinct k-mers contained by a partition determines the peak memory. Following this reasoning, we define a bin’s *load* to be the number of distinct *k*-mers in it, and will measure the *maximum bin load*, namely the highest load of any bin, as a criterion for peak memory usage.

## 3 Methods

MSP uses a minimizers scheme with a lexicographic order [10]. We denote this method **`Lexico_MSP`**. We modified MSP to employ a minimizers scheme with a UHS-based order and denote this algorithm **`UHS_MSP`**. Previous studies have shown that *k*-mers from a small UHS are more evenly distributed along the genome than lexicographic or random minimizers [15]. Hence, we reasoned that using a small UHS in the MSP algorithm would lead to a flatter distribution of bin sizes and thus reduce memory usage and runtime.

Since a pseudo-random order was shown to have better properties than lexicographic order when used in a minimizers scheme [12], we also tested a variant where the lexicographic order of the minimizers scheme in the original MSP method is replaced by a pseudo-random order. We denote this variant **`Random_MSP`**.

**`UHS_MSP`** receives as input a set of reads and generates a corresponding de Bruijn graph by the following steps. A pseudo-code of the algorithm can be found in Algorithm 1.

**Algorithm 1.**
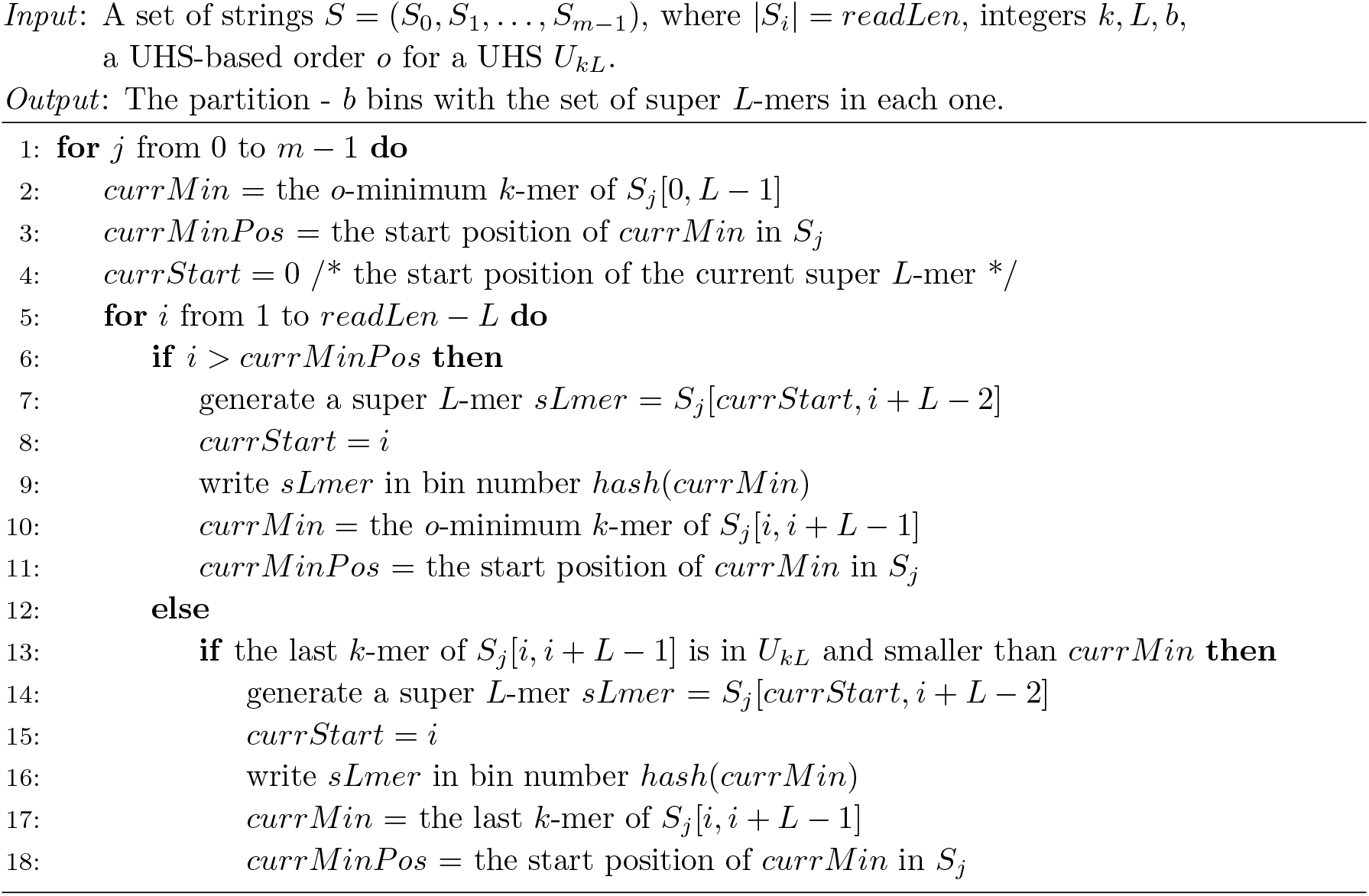
UHS Minimum Substring Partitioning.

1. **Partitioning.** This step uses a pre-generated UHS *U_kL_*. By default, we used a UHS generated by the DOCKS algorithm [15] with *k* = 12 and *L* = 60. We saved *U_kL_* in a compressed |Σ|^*k*^-long bit array with the values ‘1’ for the *k*-mers that are in *U*_*kL*_ and ‘0’ otherwise. A new order based on *U_kL_* is defined as follows: all *k*-mers in *U_kL_* are smaller than *k*-mers not in *U_kL_*, and the order of *k*-mers in *U_kL_* is random. By the definition of a UHS, a minimizers scheme based on this order selects only *k*-mers from *U_kL_* as minimizers for any *L*-long window, so the order of *k*-mers not in *U_kL_* is immaterial. We call such an order a *UHS-based minimizer order*. Reads are broken into segments (super *L*-mers) that are placed in bins as follows. For each read, all *L*-long windows are scanned and their minimizers are found. The minimizer of the currently scanned window is denoted as *currMin* and its start position is denoted as *currMinPos*. The scanning is done by sliding an *L*-long window to the right one symbol at a time, until the end of the read. After each slide, **`UHS_MSP`** checks whether *currMinPos* is still within the range of the current window. If not, it re-scans the window to find the current minimizer and updates *currMin* and *currMinPos*. Otherwise, it tests whether the last *k*-mer in the current window is smaller than *currMin* based on the UHS-based minimizer order. If so, the last *k*-mer is set as *currMin* and its start position as *currMinPos*. To enable fast comparison of *k*-mers in *U_kL_*, the pseudo-random order is implemented using a 2*k*-long bit vector *x* (the *seed*), with bits selected independently and equiprobably to be 0 or 1. For *m* ∈ *U_kL_* define *β*(*m*) = *b*(*m*) ⊕ *x*, where *b*(*m*) is the binary representation of *k*-mer *m* and “⊕” is the bit-wise xor operation. The order *o* of *m* is defined as the number whose binary representation is *β*(*m*). Hence, deciding if *o*(*m*) < *o*(*m’*) is done by two xor operations and one comparison. Each time a new minimizer is selected, a super *L*-mer is generated by merging all the *L*-long windows sharing the previous minimizer, and the label of that super *L*-mer is its minimizer (Figure 2). To obtain the prescribed number *b* of bins, a hash function is used to map the labels to a space of size *b*. A unique ID is assigned to each *L*-mer when scanning the reads. As a result, identical *L*-mers in different positions in the data are assigned different IDs. Those will be merged in the next step.
2. **Mapping and merging.** These steps are the same as in [10]. We briefly outline them here for completeness, since the changes we introduce in the partitioning step affect their efficiency. In the mapping step, each bin is loaded separately into the memory, and identical *L*-mers in different positions in the bin are combined to have the same unique integer vertex ID by generating an ID replacement table per bin. Since we expected the change in the partitioning step to create bins with sizes that are more uniformly distributed, we reasoned that the maximum bin size and the maximum memory would decrease as well. The merging step merges the ID replacement tables of all bins and generates a global ID replacement table.

The algorithm outputs sequences of IDs. Each ID is a vertex in the graph (*L*-mer) and two adjacent IDs represent an edge in the graph. This way, each read is represented by a sequence of the consecutive vertices of its *L*-mers in the graph, while identical *L*-mers have the same ID. Note that the final assembled graphs of **`Lexico_MSP`**,**`Random_MSP`** and **`UHS_MSP`** are identical. The differences in time and space performance are due to the particular schemes, which produce different bin size and load distributions.

## 4 Results

We compared **`UHS_MSP`** to the original MSP method (called here **`Lexico_MSP`**) and to MSP with random *k*-mer order (**`Random_MSP`**) in terms of speed, memory usage, particular density and maximum load on four real-life datasets (Table 1). **`UHS_MSP`** was used with the seed-based random ordering within the UHS. (Use of UHS with lexicographic ordering produced inferior results and was not tested further - see supplemental table S.1). The human chr14 and bee datasets were downloaded from the GAGE database (gage.cbcb.umd.edu/data/index.html). The *E. coli* (PRJNA431139) and the human genome data (SRX016231) were downloaded from SRA. (The bee dataset was also used in [10], but the other datasets used in that study were unavailable.) We used the same parameters as in [10] for comparison, i.e., *k* = 12, *L* = 60, and *b* = 1000. All the experiments were measured on Intel(R) Xeon(R) CPU E5-2699 v4 @ 2.20GHz server with 44 cores and 792 GB of RAM. To exclude the impact of parallelization, all measurements were done on a single core.

**Table 1:** Characteristics of the four benchmark datasets and particular density results.

|  |  |  | Particular density |  |  |
| --- | --- | --- | --- | --- | --- |
| Dataset | Size (GB) | Avg. read length | Lexico_MSP | Random_MSP | UHS_MSP |
| *E. coli* | 2.9 | 101 | 0.041 | 0.034 | **0.031** |
| Human chr14 | 9.4 | 101 | 0.073 | 0.065 | **0.062** |
| Bee | 93.8 | 124 | 0.059 | 0.056 | **0.053** |
| Human | 432 | 100 | 0.057 | 0.056 | **0.055** |

On the human data set, only **`UHS_MSP`** terminated successfully and passed all the three stages without failing. Thus, only partial results are available and reported for the other two algorithms. We note that similar problems with running MSP were reported in [11].

### 4.1 Particular density comparison

We calculated the particular density of the MSP algorithms on the four datasets by counting the number of selected positions (unique *minPos* in the partitioning step of the algorithm) and dividing it by the number of all possible positions. **`UHS_MSP`** achieved lower density than **`Lexcio_MSP`** and **`Random_MSP`** on all four datasets (Table 1), in accordance with the results of Marçias *et al*. on other genomes [12]. **`Lexico_MSP`** had the highest density. This result reaffirms the potential of **`UHS_MSP`** to achieve reduced memory usage and faster runtimes compared to the other two algorithms.

### 4.2 Performance comparison: runtime, memory and load

We compared the three algorithms in terms of three main performance criteria: (1) Runtime - the total CPU time (user time + system time); (2) Maximum memory - the maximum amount of cache memory the method used; and (3) Maximum load.

Figure 3a presents the runtime of the three algorithms in seconds per GB of input data. On all datasets where comparison was possible, **`UHS_MSP`** was the fastest. Figure 3b displays the maximum memory used by each algorithm. **`UHS_MSP`** used substantially less memory than **`Lexico_MSP`**, and achieved comparable results to **`Random_MSP`**. **`UHS_MSP`** also had the lowest maximum load on all datasets. (Figure 3c). The results show that **`UHS_MSP`** achieved a substantial improvement over the other algorithms in all three main aspects. We also measured two additional criteria: the largest bin size and the total disk space. The results for these criteria are included in supplemental figure S.1. They present consistent advantage to **`UHS_MSP`** over both **`Random_MSP`** and **`Lexico_MSP`**.

**Figure 3:**
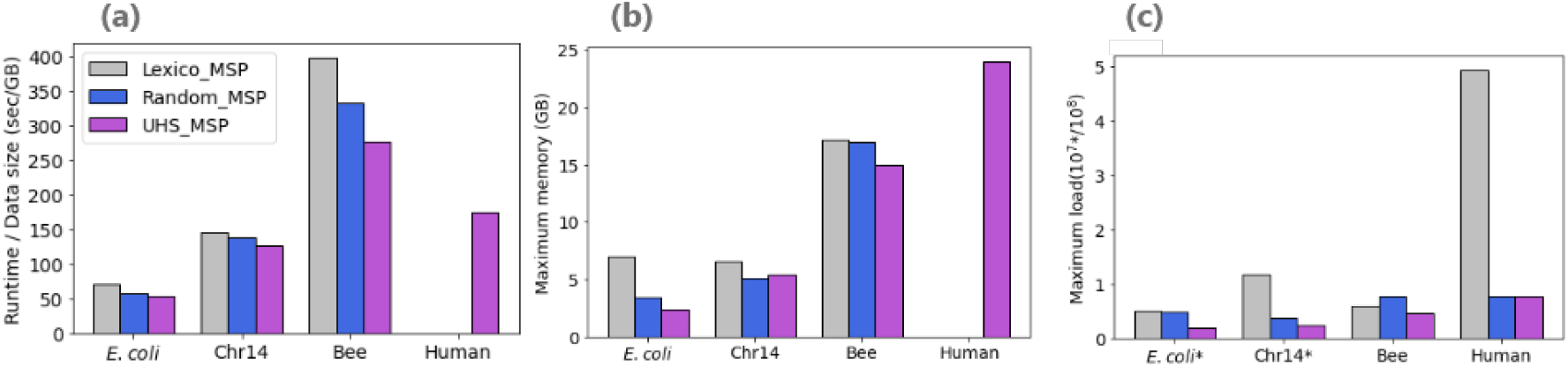
Tested metrics performance. (a) Runtime in seconds per GB of input data. (b) Maximum memory usage in GB. (c) Numbers are in 100 MB for *E. coli* and chr14 datasets, and in GB for the bee dataset. For the human dataset, the original and randomized MSP did not terminate, so runtimes and memory are not available. The reported load results are based on the partitioning and mapping steps only.

As an additional test of the robustness of **`UHS_MSP`**, we wished to gauge the effect of the pseudorandom order of the *k*-mers on the results. We ran **`Random_MSP`** and **`UHS_MSP`**,which use randomized orders, on the four datasets with five different seeds, corresponding to different pseudo-random orders (Section 3). The results are presented in Table 2. While both algorithms showed substantial performance variance across orders, overall, the results are in line with those presented in Figure 3.

**Table 2:**
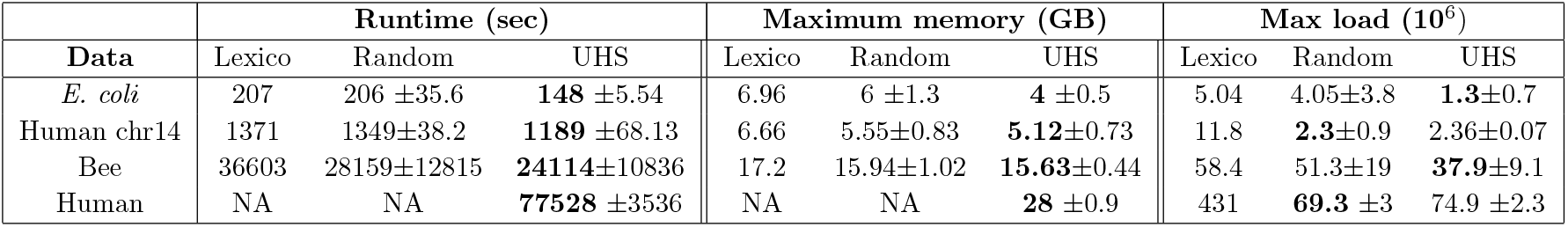
Performance across different pseudo-random *k*-mer orders. Average and standard deviation over five runs with different seeds are shown for the two algorithms that use a random order, alongside the original MSP results. On the human dataset, the original MSP as well as its pseudo-random order version did not terminate successfully.

### 4.3 The effect of parameters *k*, *L* and *b*

We tested the three algorithms on the bee data in a range of values for the parameters *k*, *L* and *b*. In each run, we kept two of the three parameters at their default values and varied the third. The results are summarized in Figure 4. Changing the number of bins shows consistent advantage to **`UHS_MSP`**, with a tendency to improve as the number of bins increases (a-c). Changing *L* shows a similar advantage to **`UHS_MSP`** (d-f). Changing *k* has a less consistent effect (g-i). Supplemental figure S.2 shows the effect of changing each of the three parameters on the largest bin size. Again, **`UHS_MSP`** was consistently better across the tested values of *L* and *b*, and less so when changing *k*.

**Figure 4:**
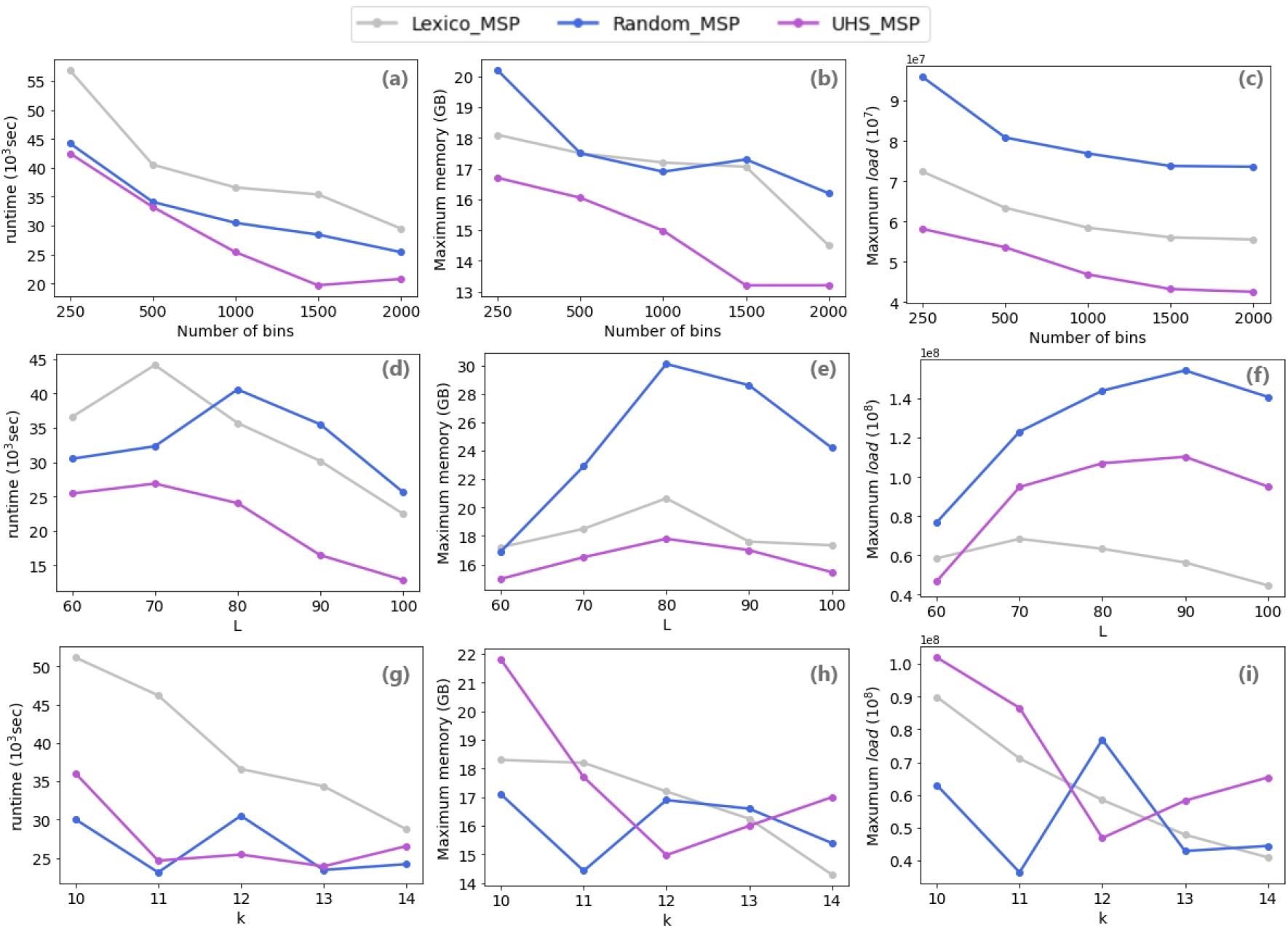
The effect of changing the number of bins, the window size and the *k*-mer size on performance. Results are for the bee dataset. (a-c) Effect of the number of bins. (d-f) Effect of the window size *L*. (g-i) Effect of *k*. For each parameter, the runtime, the maximum memory and the maximum load are shown.

Li *et al.* also tested the impact of changing *k* and *L* on **`Lexico_MSP`** [10]. The impact of varying *k* was consistent with what we observed here for that algorithm Figure 4(g-i) with reduction of resources needed as *k* increases. They also reported a similar reduction when *L* increases, unlike a less consistent picture observed here (d-f). Note however that they tested the range *L* = 31 − 63 while we tested a broader range of higher values *L* = 60 − 120. While varying the number of bins was not tested before, our tests here (a-c) show improved performance with increasing *b*, and a similar trend (with leveling off at high values) by the two other algorithms as well.

### 4.4 Resource usage in each step of the algorithm

To appreciate where the saving is achieved, we measured the resources consumed by each of the three MSP steps: partitioning, mapping and merging. Figure 5 summarizes the results on the bee dataset. The mapping step required most memory, taking an order of magnitude more memory than the partitioning and 3-5 fold more than the merging step. In all three algorithms, the mapping step was also the most time-demanding one, taking on average 67% of the time. The merging step required 20% of the time, on average, and the partitioning step was the least demanding one, taking 13% of the time. Remarkably, even though we explicitly changed only the partitioning step of the algorithm, that change led to substantial reduction in the time and memory of the mapping step. The merging step was less affected. In comparison to the original MSP algorithm, **`UHS_MSP`** required 4% more time in the partitioning step, due to the extra work required for UHS-related computations, but was 20% faster in the mapping step. Note that for the sake of our tests, we did not utilize the possibility of parallelizing the mapping step. Future work can thus focus on improvements to the mapping step.

**Figure 5:**
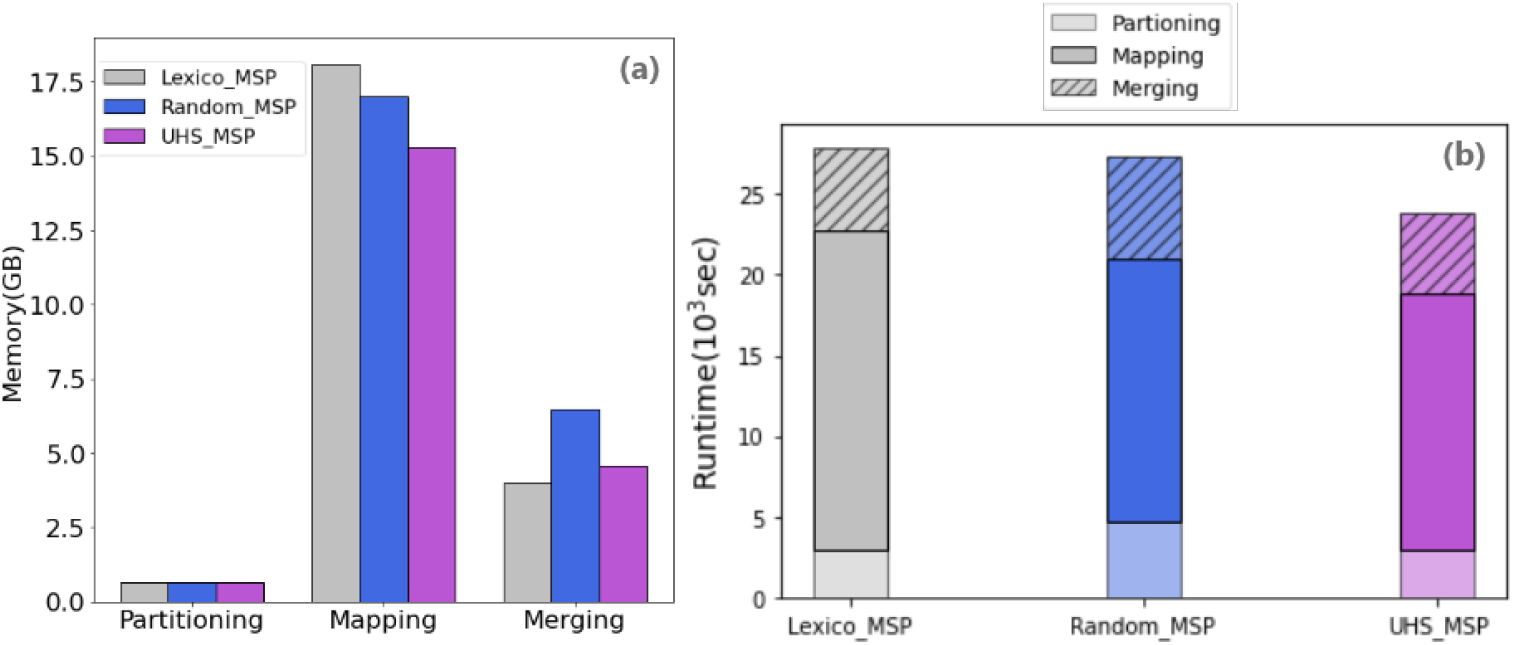
Resources taken by each algorithm and each step of the algorithm on the bee dataset. (a) Maximum Memory. (b) Runtime.

## 5 Discussion

In this study, we incorporated a UHS-based minimizers scheme in a fundamental HTS task: de Bruijn graph construction. By creating partitions based on fewer *k*-mers and with better statistical properties, we achieved speedups and reduced memory usage in genomic assembly. To the best of our knowledge, this is the first demonstration of the practical advantage of using UHSs in a genome assembly application.

Our study raises several open questions: to what extent can further improvements in the generation of smaller UHSs improve the de Bruijn graph construction? Currently the complexity of minimum size UHS still remains an open problem, though closely-related problems were shown to be computationally hard [12, 15]. Can one express the expected amount of resources needed by the **`UHS_MSP`** algorithm (and by its separate steps), as a function of the key parameters *k*, *L* and *b*? Obtaining such an estimate, even under a simple model such as the random string model [10], can guide one to optimize the combination of parameter values, which as we have seen tend to interact in a rather complex way (Figure 4). What is the relation between the largest bin size and the maximum memory usage? Can one improve the mapping of multiple minimizers into a bin? Some applications (e.g., [4]) sample the input data in order to map minimizers to bins more efficiently. Integrating such a method into MSP can improve memory and runtime performance.

Li *et al.* [10] argued that the the maximum load determines the peak RAM consumption. Our results were not consistent with this claim. (For example, see the results on the bee data in Figure 3 and Figure 4 (d-f)). Further investigation revealed that some of the incoherence is due to Java’s garbage collector. Changing Java’s limit for the heap size reduced memory consumption substantially. For example, when running the chr14 dataset with a limit of 4GB, the peak memory went down from 6.6GB to 2.7GB, with no increase in running time. In other runs even larger peak memory changes were observed depending on the limit, but at the expense of longer runtimes. Since the measured memory of a Java process is not reliable, and depends on the machine and on Java’s memory flags, an implementation in a language with explicit memory management (e.g. c++) is preferable.

In parallel to our study, a work by Nystrom-Persson *et al.* [13] also implemented UHS in a sequencing application. The authors combined UHS and frequency counts in a k-mer counting application and achieved substantial memory saving. Given the improvement now achieved by UHSs in two different sequencing applications, it is tempting to believe that practical improvements can be achieved in other applications that utilize minimizers, e.g. *BCALM* 2 [2] (which utilizes frequency-based minimizers, given the results in [13]), applications that perform partitioning of sequences as a preprocessing step for efficient parallel processing and storage [20, 3, 4, 9], sequence similarity estimation [7, 14], and others.

## 6 Acknowledgments

This study was supported in part by the Israeli Science Foundation grant 1339/2018 and grant No. 3165/19, within the Israel Precision Medicine Partnership program (to R.S.), and by German-Israeli Project DFG RE 4193/1-1. Y.B. and L.P. were partially supported by fellowships from the Edmond J. Safra Center for Bioinformatics at Tel-Aviv University. L.P. was also supported in part by postdoctoral fellowships from the Planning Budgeting Committee (PBC) of the Council for Higher Education (CHE) in Israel.

---

**Table S.1:** Performance of MSP using UHS and lexicographic ordering compared to the other three algorithms on the bee dataset. The right column shows the results of MSP using UHS and lexicographic ordering,

| Method | Lexico_MSP | Random_MSP | UHS_MSP | Lexico_UHS |
| --- | --- | --- | --- | --- |
| Runtime(sec) | 36603 | 30498 | 25443 | 40246 |
| Max memory(GB) | 17.2 | 16.9 | 14.98 | 17.7 |

**Figure S.1:**
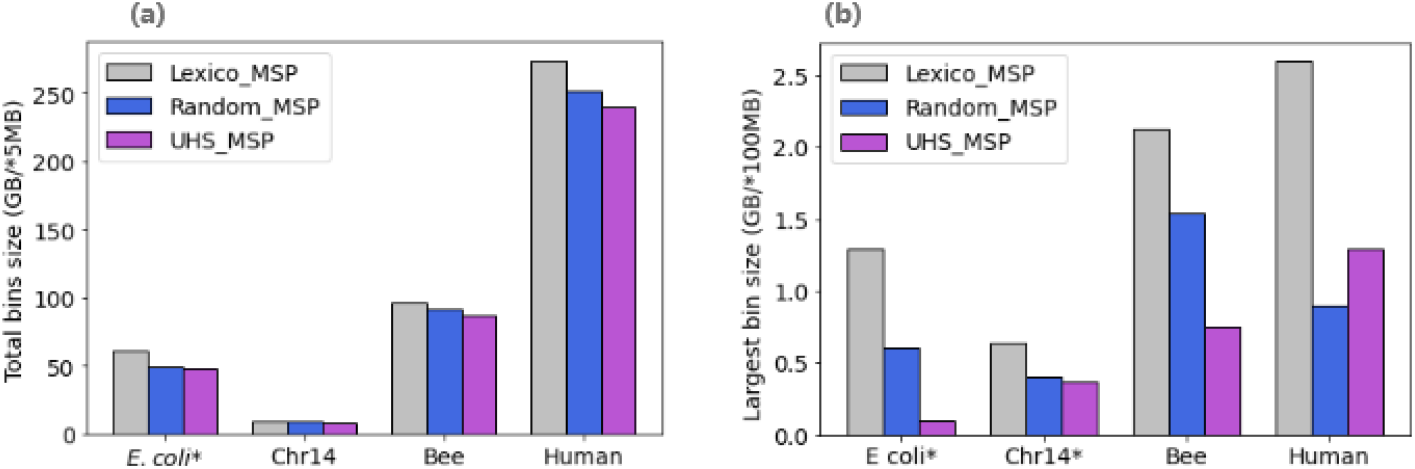
Additional tested metrics performance. (a) **Total bins size.** The total size of the bins stored on the disk is shown. Numbers are in 5MB for *E.coli* dataset and in GB for chr14, bee and human datasets. (b) **Largest bin size.** Numbers are in 100 MB for *E. coli* and chr14 datasets, and in GB for the bee and human datasets.

**Figure S.2:**
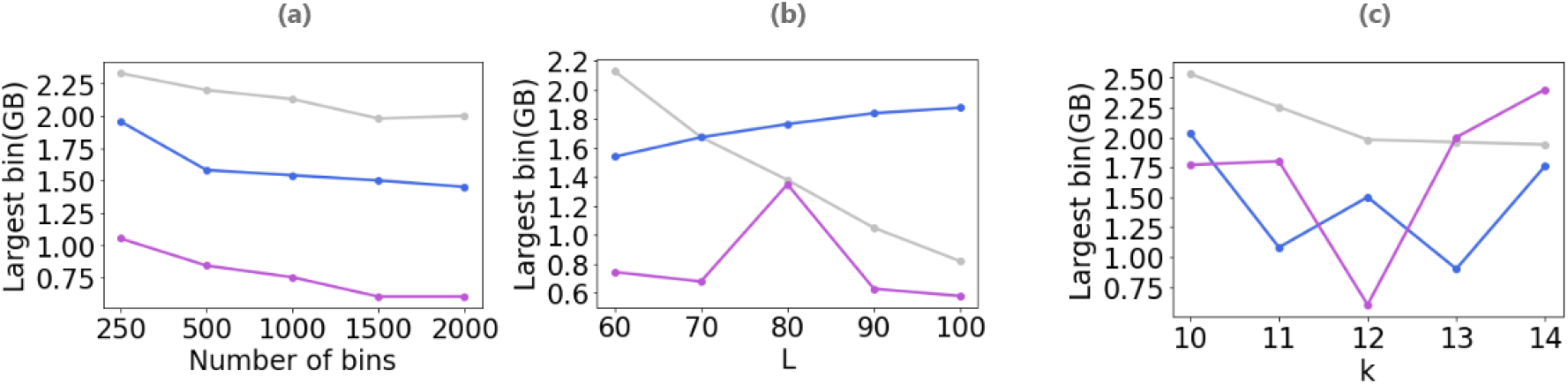
The effect of changing the number of bins, the window size and the *k*-mer size on the largest bin size. Results are for the bee dataset. (a) Effect of the number of bins. (b) Effect of the window size *L*. (c) Effect of *k*.

## References

[1] S. Anders, P. T. Pyl, and W. Huber. HTSeq—a Python framework to work with high-throughput sequencing data. Bioinformatics, 31(2):166–169, 2015.

[2] R. Chikhi, A. Limasset, and P. Medvedev. Compacting de Bruijn graphs from sequencing data quickly and in low memory. Bioinformatics, 32(12):i201–i208, 06 2016.

[3] R. Chikhi, A. Limasset, and P. Medvedev. Compacting de Bruijn graphs from sequencing data quickly and in low memory. Bioinformatics, 32(12):i201–i208, 2016.

[4] S. Deorowicz, M. Kokot, S. Grabowski, and A. Debudaj-Grabysz. KMC 2: fast and resource-frugal k-mer counting. Bioinformatics, 31(10):1569–1576, 2015.

[5] S. Grabowski and M. Raniszewski. Sampling the suffix array with minimizers. In International Symposium on String Processing and Information Retrieval, pages 287–298. Springer, 2015.

[6] W. Huber, V. J. Carey, R. Gentleman, S. Anders, M. Carlson, B. S. Carvalho, H. C. Bravo, S. Davis, L. Gatto, T. Girke, et al. Orchestrating high-throughput genomic analysis with Bioconductor. Nature Methods, 12(2):115, 2015.

[7] C. Jain, A. Dilthey, S. Koren, S. Aluru, and A. Phillippy. A fast approximate algorithm for mapping long reads to large reference databases. In S. Sahinalp, editor, Research in Computational Molecular Biology. RECOMB 2017. Lecture Notes in Computer Science, vol 10229, pages 66–81. Springer, 2017.

[8] G. Kucherov. Evolution of biosequence search algorithms: a brief survey. Bioinformatics, 2019.

[9] Y. Li et al. MSPKmerCounter: a fast and memory efficient approach for k-mer counting. arXiv preprint arXiv:1505.06550, 2015.

[10] Y. Li, P. Kamousi, F. Han, S. Yang, X. Yan, and S. Suri. Memory efficient minimum sub-string partitioning. In Proceedings of the VLDB Endowment, volume 6, pages 169–180. VLDB Endowment, 2013.

[11] S. C. Manekar and S. R. Sathe. A benchmark study of k-mer counting methods for high-throughput sequencing. GigaScience, 7(12), 10 2018. giy125.

[12] G. Marçais, D. Pellow, D. Bork, Y. Orenstein, R. Shamir, and C. Kingsford. Improving the performance of minimizers and winnowing schemes. Bioinformatics, 33(14):i110–i117, 2017.

[13] J. Nyström-Persson, G. Keeble-Gagnère, and N. Zawad. Compact and evenly distributed k-mer binning for genomic sequences. Bioinformatics, 03 2021. btab156.

[14] B. D. Ondov, T. J. Treangen, P. Melsted, A. B. Mallonee, N. H. Bergman, S. Koren, and A. M. Phillippy. Mash: fast genome and metagenome distance estimation using MinHash. Genome Biology, 17(1):132, 2016.

[15] Y. Orenstein, D. Pellow, G. Marçais, R. Shamir, and C. Kingsford. Designing small universal k-mer hitting sets for improved analysis of high-throughput sequencing. PLoS Computational Biology, 13(10):e1005777, 2017.

[16] J. A. Reuter, D. V. Spacek, and M. P. Snyder. High-throughput sequencing technologies. Molecular Cell, 58(4):586–597, 2015.

[17] M. Roberts, W. Hayes, B. R. Hunt, S. M. Mount, and J. A. Yorke. Reducing storage requirements for biological sequence comparison. Bioinformatics, 20(18):3363–3369, 2004.

[18] M. Roberts, B. R. Hunt, J. A. Yorke, R. A. Bolanos, and A. L. Delcher. A preprocessor for shotgun assembly of large genomes. Journal of Computational Biology, 11(4):734–752, 2004.

[19] S. Schleimer, D. S. Wilkerson, and A. Aiken. Winnowing: local algorithms for document finger-printing. In Proceedings of the 2003 ACM SIGMOD International conference on Management of data, pages 76–85. ACM, 2003.

[20] D. E. Wood and S. L. Salzberg. Kraken: ultrafast metagenomic sequence classification using exact alignments. Genome Biology, 15(3):R46, 2014.

[21] C. Ye, Z. S. Ma, C. H. Cannon, M. Pop, and W. Y. Douglas. Exploiting sparseness in de novo genome assembly. In BMC Bioinformatics, volume 13, page S1. BioMed Central, 2012.

[22] H. Zheng, C. Kingsford, and G. Marçais. Lower density selection schemes via small universal hitting sets with short remaining path length. In International Conference on Research in Computational Molecular Biology, pages 202–217. Springer, 2020.

